# Pair-bond strength is consistent and related to partner responsiveness in a wild corvid

**DOI:** 10.1101/2023.12.16.571986

**Authors:** Rebecca Hooper, Luca G. Hahn, Guillam E. McIvor, Alex Thornton

## Abstract

The need to maintain strong social bonds is widely held to be a key driver of cognitive evolution. This assumes that the maintenance of strong bonds is a stable trait that is cognitively demanding but generates fitness benefits, and so can come under selection. However, these fundamental micro-evolutionary tenets have yet to be tested together within a single study system. Combining observational and experimental behavioural data with long-term breeding records, we tested four key assumptions in wild jackdaws (*Corvus monedula*), corvids whose long-term pair-bonds exemplify the putative social drivers of cognitive evolution in birds. We found support for three assumptions: (1) pair-bond strength varies across the population, (2) is consistent within pairs over time and (3) is positively associated with a measure of socio-cognitive performance. However, we did not find evidence that stronger pair-bonds lead to better fitness outcomes (prediction 4). While strongly bonded pairs were better able to adjust hatching synchrony to environmental conditions, they did not fledge more or higher quality offspring. Together, these findings provide important evidence that the maintenance of strong pair bonds is linked to socio-cognitive performance and facilitates effective coordination between partners. However, they also imply that these benefits may not be sufficient to explain how selection acts on social cognition. We argue that evaluating how animals navigate trade-offs between investing in long-term relationships versus optimising interactions in their wider social networks will be a crucial avenue for future research.

## Introduction

Many social animals exhibit differentiated social relationships or bonds with others, which can be defined operationally as repeated affiliative interactions between individuals in close proximity (1). These bonds can enable the exchange of behavioural commodities such as social support (2), food (3), information (4) and affiliation (5), and may generate important fitness benefits (6,7). For instance, numerous studies across different taxonomic groups indicate that individuals that form strong social bonds and are well integrated within social networks show elevated survival or reproductive success (6,7). The formation and maintenance of these strong bonds may be facilitated by abilities to track and respond to partners’ behaviour and make strategic social decisions (8–10). Thus, the advantages of being able to establish strong social bonds could, in principle, generate selection on information-processing, or cognitive abilities (9,11).

These ideas are encapsulated by the highly influential but controversial *Social Intelligence (or Social Brain) Hypothesis* (*SIH* or *SBH*) (12–16); for critiques see (17–26). While variants of the hypothesis differ in their emphasis (e.g., on “Machiavellian” manipulations (14), social learning (27) or cooperation (28)), they share a common focus on the importance of navigating social relationships as a central driver of cognitive and brain evolution. For example, in primates, individuals often form multiple stable relationships (29). Accordingly, there is evidence that primate species that live in bigger groups (with a greater number of potential social partners) tend to have bigger brains (15,17,30), but see (26). The fundamental logic of the *SIH* may be extended to other taxa. For instance, like primates, some birds – notably corvids and parrots - are highly social and are also renowned for their large brains and sophisiticated cognitive abilities (31,32). However, avian brain size appears to be associated not with the quantity of social connections (21,25,33,34) but with long-term pair-bonding (25,35) (though again this finding is controversial (21)). Some authors therefore argue that pair-bonding imposes selective pressure on cognition. This bird-focused version of the *SIH* (sometimes known as the *Relationship Intelligence Hypothesis* (25) posits that in long-term pair-bonds with interdependent fitness outcomes, partners must track information about one another to minimise conflict and enable effective cooperation, and that this is both cognitively demanding and leads to increased reproductive success. The benefits of such “relationship intelligence” are therefore suggested to be a key driver of cognitive evolution in birds (25,35).

To date, investigations of the *SIH* have relied largely on comparative analyses of neuroanatomy and behaviour, often with contentious and contradictory results (19,23,26). There is growing recognition that if we are to understand the potential for selection to act on cognitive traits, micro-evolutionary approaches focusing on the causes and consequences of within-species variation are also necessary (18,36,37). If bond strength varies within populations but is consistent within individuals and dyads, and leads to increased fitness, the ability to form and maintain strong social bonds could come under positive selection. In birds, there is evidence that pair-bond strength – the degree of affiliative behaviour that mating partners engage in – varies between pairs in captive (38–42) and wild (43) populations. Evidence for consistency of pair-bond strength is limited to between-year repeatability of spatial proximity in wild greylag geese (*Anser anser*) (43) and within-year stability of affiliative ‘clumping’ in captive zebra finches (*Taeniopygia guttata*) (42). Studies also suggest that the duration of pair bonds is positively linked to reproductive success (e.g., (44)). For instance, captive cockatiels (*Nymphicus hollandicus*) with stronger bonds have increased fledging success than those with weaker bonds (45), and captive zebra finches with more stable bonds start their breeding attempts before those with less stable bonds (42). However, to understand whether and how selection acts on relationship strength in natural contexts, fitness outcomes must be investigated in wild populations. Moreover, despite being a foundational assumption of the *SIH* (9), the link between social bonding and cognition remains unclear. Indeed, in principle, interacting repeatedly with the same partner(s) could reduce uncertainty and allow partners to pool their skills, thus reducing cognitive demands (46,47). Conversely, information-processing abilities that enable individuals to detect and respond to a partner’s state could facilitate the maintenance of successful cooperative relationships (25,48). To evaluate these possibilities, an important step is to examine whether individual socio-cognitive performance is positively associated with the maintenance of strong social bonds.

Here, we tested four key assumptions of the *SIH* within one study system, wild jackdaws (*Corvus monedula*), a highly social corvid species. Like other corvids, jackdaws have large brains and exhibit social behaviours including social information use (8,49), social support (25), and cooperation (8,50). Crucially, jackdaw societies are centred around long-term, genetically monogamous pair-bonds (51–55) - the key putative social drivers of cognitive evolution in birds (25,35). As cavity nesters, they also take readily to nest boxes, allowing the monitoring of reproductive success. We predicted, first, that pair-bond strength should be variable between pairs and, second, that pair-bond strength should be consistent within pairs, i.e., repeatable between observations both within and across years. Third, if cognition and bond-strength are linked, we predicted that individuals that have stronger pair-bonds should be more responsive to their partner’s affective state. To test this, we used measures of responsiveness from an experiment where females were exposed to a stressor in the absence of their male partner, such that males’ responses reflect their ability to detect and respond to information about their partner’s distress (56). Finally, we predicted that variation in pair-bond strength should be linked to the pair’s ability to respond to changing environmental conditions and, ultimately, to their reproductive success. Specifically, we expected that pairs with stronger bonds would (a) be better able to adjust breeding attempts to current conditions through changes in hatching asynchrony and (b) successfully rear more fledglings, with greater overall fledgling mass.

## Methods

### Subjects and study sites

Behavioural, morphometric, and breeding data were collected from colour-ringed jackdaws during the breeding seasons in 2014, 2015, 2018, and 2019 at three nest box colonies in Cornwall, UK. The study sites were the University of Exeter’s Penryn campus (Site X: 50°10’23"N, 5°7’6"), a churchyard and its adjacent fields in Stithians (Site Y: 50°11′26″N, 5°10′51″W), and a private farm near Stithians (Site Z: 50°11′56″N, 5°10′9″W). In this study, data from 125 individuals (63 females, 62 males) were included. Further details of the study population, including breeding data, and capture and ringing methods, are given in the Supplementary Material.

### Video data

During the breeding season, lasting from March to June, both members of the pair cooperate to build the nest (50,53), and the female incubates the eggs while her partner brings her food (53). We captured video footage of jackdaws inside their nest box during the nest-building and incubation stages of the breeding seasons. We recorded footage in the early morning, starting shortly after sunrise, using hidden CCTV cameras placed inside nest boxes. The footage was coded with a detailed behavioural ethogram to allow fine-scale quantification and analysis of pair-bond strength (Table S1). In the nest-building stage, we recorded 142.18 hours of footage across 54 videos (mean length = 2.63 ± 1.13 hours per video) for 39 pairs. In the egg laying/incubation stage, we recorded 362.24 hours of footage across 132 videos (mean length = 2.74 ± 1.07 hours per video) for 65 pairs.

### Data processing and statistical analysis

All data processing and statistical analysis was conducted in R version v4.0.2 (57).

#### Quantifying pair-bond strength

Behaviours recorded from internal nest box video were standardised by the length of the video. To quantify pair-bond strength (Figure S1) and identify interrelationships between potential affiliative behaviours (41,58,59), we used principal components analysis (PCA) (package: *psych* (60)). For the PCA, we considered the following behaviours previously hypothesised to be important affiliative behaviours between bird partners: ‘food-share’ (3); ‘contact’ (52); ‘allopreen’ (5), ‘time together’ (39), and ‘copulation’ (41). We also included ‘male visit rate’ in the incubation stage because females must remain in the nest box to incubate, but the rate at which the males visit their partner may vary and may be correlated with other affiliative behaviours. Finally, we included ‘chatter’, a distinctive call that partners often make when together at the nest box. Allopreening was split into ‘male-initiated’ and ‘female-initiated’ for the nest-build stage, but not at the incubation stage because almost all allopreening (94.55%) was male-initiated. The first principal component, PC1, explained a substantial proportion of variation in the data for both the nest-building (43.3%: dominated by allopreening, contact and time together; Figure S2a; Table S2) and incubation stage (59.5%: dominated by male chatter, female chatter, allopreening, contact and time together; Figure S2b; Table S2). We therefore used each pair’s PC1 value as the measure of ‘pair-bond strength’ in both instances when constructing statistical models.

#### Statistical modelling

For cases where we built competing models (see Supplementary Material, Methods), we compared models using Akaike’s Information Criterion (AIC) (61). If models differed in AIC by 2 or more, we selected the model with the lowest AIC. Otherwise, the model with the best diagnostic plots was retained (assessed using the package *DHARMa* (62)). If diagnostics for the full models were essentially equivalent, we examined model diagnostics for models once influential points were excluded, and the best fitting model retained. We identified influential points as any datapoints more than four times the mean Cook’s distance. We ran each final model both with and without influential points. Models without influential points are presented in the main manuscript (and corresponding tables and figures). If results from full models and models without influential points differed, we report the results of both models. We tested all models for zero-inflation and dispersion. No final models were over-dispersed or zero-inflated and all showed acceptable model fit.

### Does pair-bond strength (1) vary between pairs and is it (2) consistent within pairs?

We tested whether within-pair-bond strength within pairs remained consistent over time (both within and across years), using repeatability analysis in the package *rptR* (63). Specifically, we tested whether pair-bond strength was repeatable within-year and between-years for both the nest-building and incubation stage. We first ran repeatability models with no covariates to obtain an unadjusted estimate of repeatability of pair-bond strength. Following this, we controlled for covariates and obtained an adjusted repeatability estimate. Covariates for the within-year models were days since the female’s fertile period and time the video was started. Given within-year repeatability, a mean value of pair-bond strength per year was calculated, and we used this value to estimate repeatability between years. For between-year models, we included the age of the male and year as covariates. We did not include days since the female’s fertile window and video start time as covariates due to the use of mean value of within-year pair-bond strength as the response variable. In all models, we log-transformed pair-bond strength. For each model, parametric bootstrapping (n_boot_ = 1000) quantified uncertainty while significance testing was implemented using likelihood ratio tests (LRT) and through the permutation of residuals (n_perm_ = 1000).

### 3) Is pair-bond strength related to partner responsiveness?

In 2019, we ran a field experiment to test whether males responded to their partner’s distress (56). To do this, we used playbacks to expose incubating females (N = 27) to the sound of a foreign male at the nest box – an important stressor as foreign males subject nesting females to violent attacks. Focal females’ male pair-bonded partners were absent from the area during playbacks, and therefore blind to the stressor. We supplemented experimental data with data from natural forced extra-pair copulation events where we had a measure of male behaviour towards his female partner both pre- and post-stressor, and the male was absent for the stressor (N = 6; experimental and natural data produced qualitatively the same results). In this original study we found that, overall, males altered their behaviour after their partner had experienced the stressor, indicating that they responded to their partner’s state. However, the nature and magnitude of changes in male behaviour was highly variable ((56); Figure 2). While the overall effect was that males reduced visit rates and affiliation (suggesting males use indicators of female state to minimise their own exposure to risk), some males showed the opposite effect, and a few did not change their behaviour at all. Here, we therefore measured ‘partner responsiveness’ as the absolute change in male-initiated direct affiliative behaviour (contact and allopreening) towards his partner for 1.5 hours pre- and 1.5 hours post-stressor (the time period over which the strongest effects were found in the original study). In our model, partner responsiveness, the dependent variable, was log-transformed for improved model fit. We included pair-bond strength, data type (experimental or natural), whether the male returned before the female and minimum age of the male (which resulted in a better model fit than minimum number of years together) as covariates for the full model. Due to convergence issues, minimum age of the male was not included as a predictor in the model without influential points; however, the relationship between pair-bond strength and partner responsiveness was qualitatively identical for both models. We followed this analysis with two further models to test: (1) whether female behaviour changed as a function of pair-bond strength, and thus whether males could simply have been responding to the differential magnitude of female behavioural change and (2) whether males were responding to overt changes in female behaviour, we also tested whether males responded more strongly when females showed a greater magnitude of behavioural change in response to the stressor.

### (4) Do pairs with stronger bonds have higher reproductive success?

We built GLMMs to test whether pair-bond strength was linked to fitness outcomes. Because pair-bond strength was repeatable during the incubation stage of the breeding season (see Results), we tested whether pair-bond strength measured in this stage was associated with reproductive outcomes. For each response variable reflecting one aspect of reproductive success (see below) we tested competing models which included either ‘minimum number of years together’ or ‘minimum male age’. We always included ‘pair-bond strength’ (per video), ‘male tarsus’, ‘female tarsus’, ‘lay date’, ‘year’, and the ‘rate of male provisioning during female incubation’ (uncorrelated with affiliative behaviours yet potentially important for reproductive success) as fixed effects. Furthermore, we included ‘pair ID’ and ‘site’ as random effects. For all fitness outcomes, we compared models with linear and quadratic pair-bond strength, to test for directional and stabilising selection on pair-bond strength (64). We also compared models with and without an interaction term between pair-bond strength and year. We tested this interaction because how selection acts on behaviour can vary according to environmental conditions (e.g. (65)).

#### (a) Coordination of hatching synchrony

Daily observation allowed us to monitor the date of egg-hatching in order to calculate hatching synchrony. Hatching synchrony was calculated as the date of the last hatch minus the date of the first hatch, divided by the number of eggs that hatched. We examined whether a strong pair-bond may facilitate fitness-enhancing responsiveness to changing environmental conditions across breeding seasons by testing the relationship between hatching synchrony as a response variable and an interaction between year and pair-bond strength. More asynchronous hatching is thought to be more advantageous in resource-poor years because the brood is quickly reduced, thus increasing the probability that a small number of chicks survive rather than entire brood failure (66,67), but see (68). Conversely, more synchronous hatching is thought to be advantageous in resource-rich years (66). In jackdaws, females incubate their eggs while being provisioned by their partner (69). The synchronicity of hatching depends on the female’s incubation behaviour (70), which in turn is influenced by the male’s provisioning behaviour (69). Hatching synchrony may therefore be related to how well a pair are able to coordinate their behaviour in the face of environmental variability.

#### (b) Success in fledging offspring

The measures for fledging success we tested for each pair (per year) were (1) number of fledglings, (2) total mass of fledglings, and (3) proportion of hatched chicks that fledged. Because of low levels of variance in number of fledglings per year (72.87% of pairs fledged two or three offspring), we also tested the cumulative number of fledglings per pair over five years. This required sub-setting the data to pairs for whom we had five years of reproductive success data (N = 12). For this model, pair-bond strength was calculated as the mean value of pair-bond strength per pair across all available data points in the incubation stage. To test how reproductive success varied across years for the whole population, we also examined the relationship between year and population-wide number and mass of fledglings. For these models, we included ‘year’, ‘male and female tarsus length’, ‘minimum male age’ or ‘years together’, and ‘lay date’ as predictor variables. We included ‘pair ID’ as a random effect, but site was not included due to convergence issues. These models allowed us to gain insight into whether some years appeared particularly difficult, most likely due to limited resource availability. To test whether hatching synchrony interacted with year to influence reproductive success, we ran GLMMs with (1) number of fledglings, (2) total mass of fledglings, and (3) proportion of hatched chicks that fledged across the entire population as response variables and hatching synchrony as predictor. However, models (1) and (2) would not converge with the inclusion of site, so it was removed. For each response variable, we compared model performance with and without an interaction term between year and hatching synchrony.

## Results

### Does pair-bond strength (1) vary between pairs and is it (2) consistent within-pairs?

Pair-bond strength varied considerably between pairs both during the nest-building and the incubation stage (Figure 1). Pair-bond strength was not repeatable in the nest-building stage (Supplementary Results) but during the incubation stage (for which sample sizes were higher) pair-bond strength was highly repeatable, both within-year (N = 21 pairs, adjusted R = 0.65, 95% CI[0.34, 0.87], Pperm = 0.003, PLRT = <0.01) and between-years (N = 34 pairs, adjusted R = 0.50, 95% CI[0.26, 0.71], Pperm = <0.01, PLRT <0.01; Figure 1). There was one highly affiliative pair (evident in Figure 1), but results were robust to their removal (within-year: adjusted R = 0.49, 95% CI[0.08, 0.80], Pperm = 0.03, PLRT = 0.02; between-year: adjusted R = 0.29, 95% CI[0.03, 0.57], Pperm = 0.02, PLRT = 0.05).

**Figure 1.**
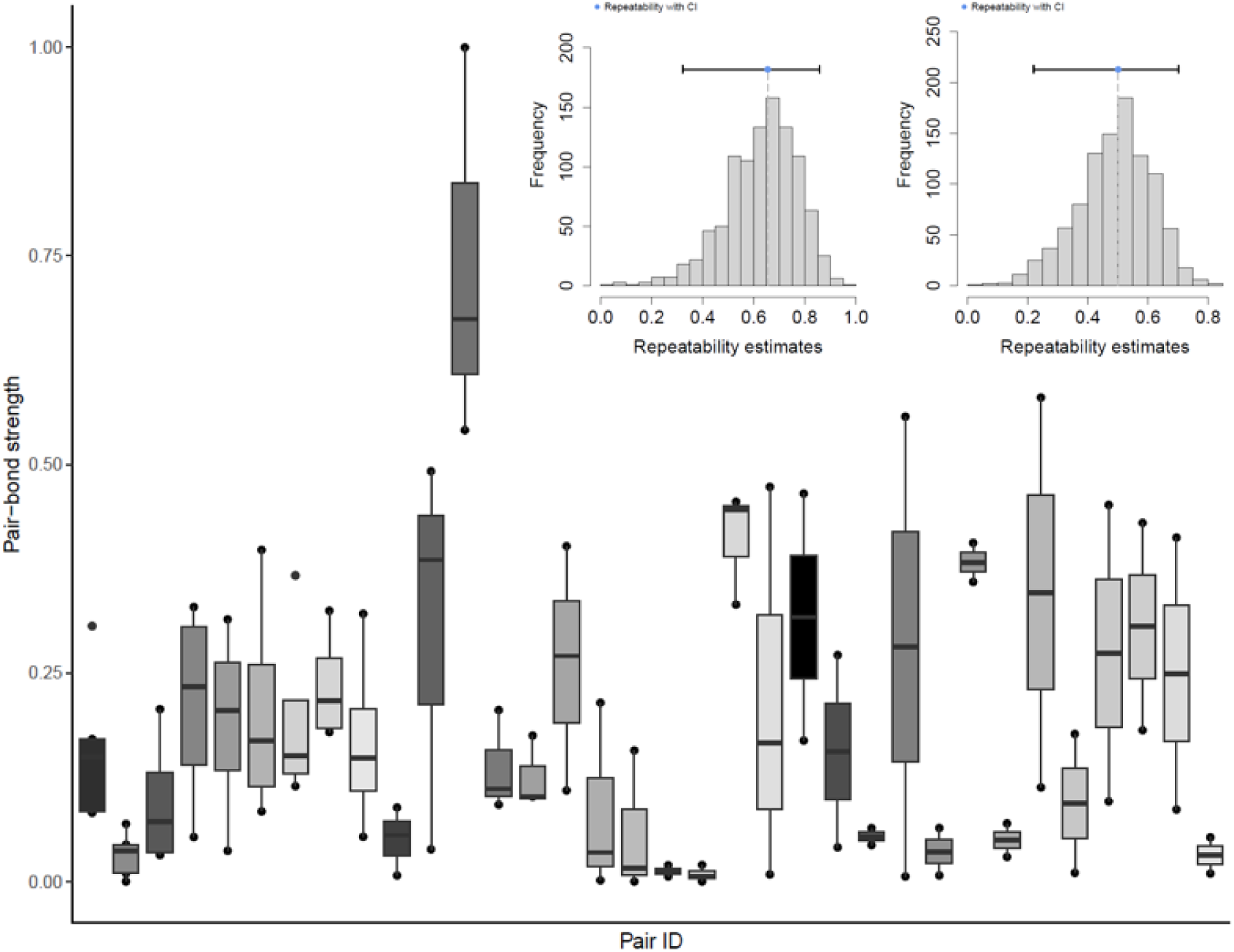
Pair-bond strength per pair, quantified during the incubation stage. Each datapoint represents the strength of a pair’s bond in a single year; thus, boxplots show the range of bond strength per pair across multiple years. Repeatability plots are depicted in the top right corner, displaying estimates (including confidence intervals shown as horizontal bars) of within-year bootstrap repeatability (left), and between-year bootstrap repeatability (right). Jackdaw pair-bond strength was significantly repeatable both within and between years, with confidence intervals not overlapping with zero, and the results were robust to the removal of a pair with extraordinarily high pair bond strength. Note that for visualisation purposes, pair-bond strength (log-transformed PCA values) is standardised between 0 and 1.

**Figure 2.**
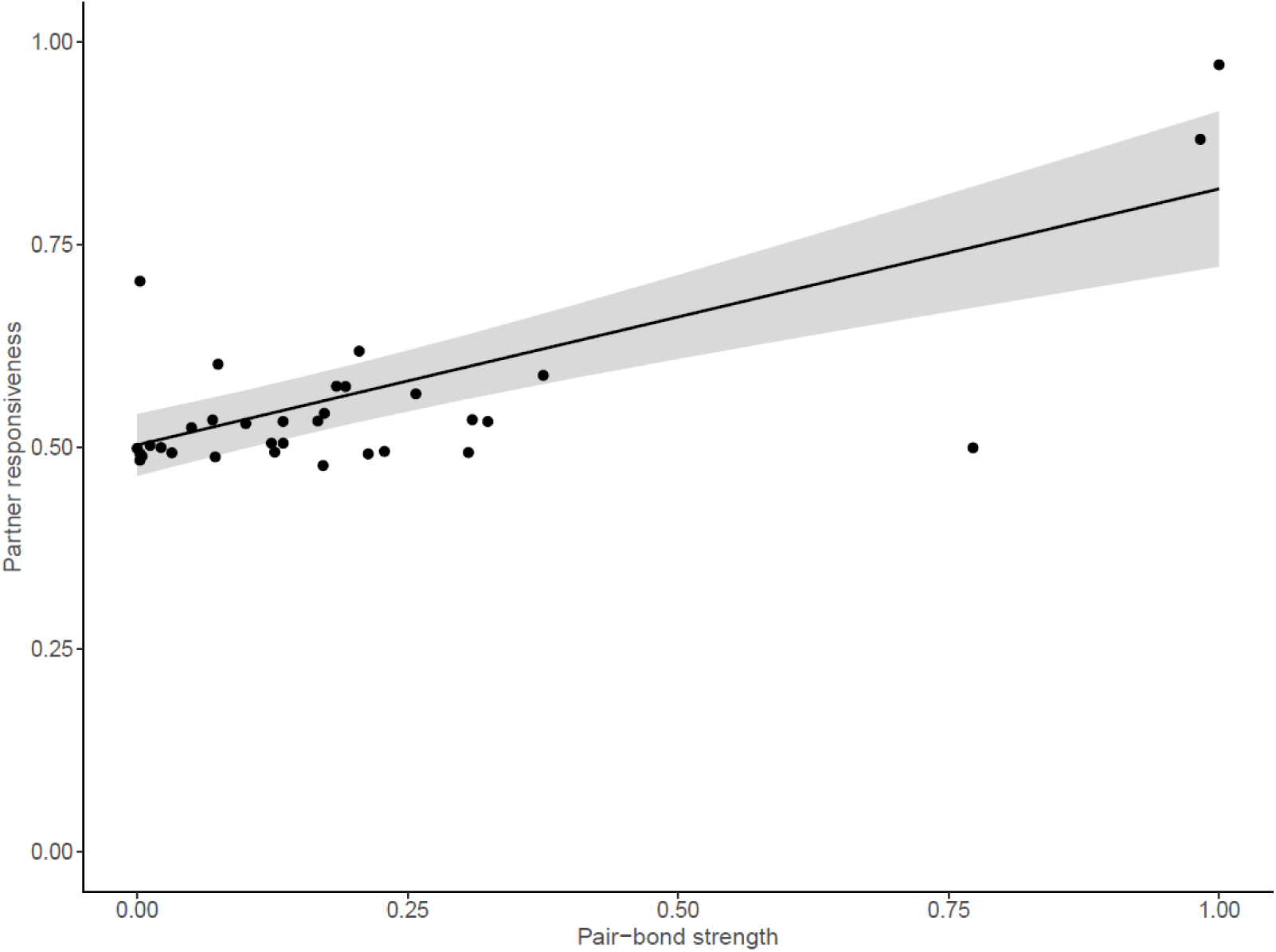
Pair-bond strength was significantly positively correlated with partner responsiveness (a measure of socio-cognitive performance in experimental tests), both in the full model and with influential points excluded. Note that for visualisation purposes, pair-bond strength and partner responsiveness are standardised between 0 and 1.

### (3) Is pair-bond strength related to partner responsiveness?

There was a significant relationship between pair-bond strength and the responsiveness of the male to his partner’s distress (Figure 2), where males in stronger pair-bonds showed a larger absolute change in behaviour following their partner experiencing a stressor (N pairs = 33; β = 0.10, SE = 0.01, 95% CI [0.07, 0.13], P < 0.001). This result was robust to the removal of influential points (N pairs = 30; β = 0.12, SE = 0.01, χ^2^ = 137.97, 95% CI [0.10, 0.14], P < 0.01). Previous analyses have reported weak evidence for slight declines in female chatter calls and begging rates post-stressor (56). However, male responses were not linked to any detectable change in female behaviour: There was no significant relationship between the absolute change in female begging or chatter and pair-bond strength (begging rate: N pairs = 24, β = −0.14, SE = 0.09, χ^2^ = 2.65, 95% CI [-0.31, 0.03], P = 0.10; chatter duration: N pairs = 23, β = −0.15, SE = 0.15, χ^2^ = 0.95, 95% CI [-0.45, 0.15], P = 0.33), or the magnitude of the female’s change in vocalisations and the magnitude of the male’s behavioural change (begging rate: N pairs = 23, β = 0.19, SE = 01.88, χ^2^ = 0.01, 95% CI [-3.50, 3.88], P = 0.92; chatter: N pairs = 24, β = −0.56, SE = 0.86, Х2 = 0.42, 95% CI [-2.25, 1.13], P = 0.52).

### (4) Do pairs with stronger bonds have higher reproductive success?

#### (a) Coordination of hatching synchrony

The interaction between pair-bond strength and year was significantly associated with hatching synchrony (full model: N datapoints = 102, N pairs = 50, χ^2^ = 8.00, p = 0.046; Figure 3), although the model term only bordered significance with influential points removed (N datapoints = 92, N pairs = 47, χ^2^ = 7.06, P = 0.07). Pairwise comparisons between years were consistent for the full model (2018 relative to 2015: β = 0.11, SE = 0.05, 95% CI [0.01, 0.21], p = 0.02) and the model with influential points removed (β = 0.22, SE = 0.10, 95% CI [0.03, 0.42], P = 0.03). The full model also revealed a significant difference between 2018 and 2014 (β = 0.22, SE = 0.08, 95% CI [0.06, 0.38], P < 0.01).

**Figure 3.**
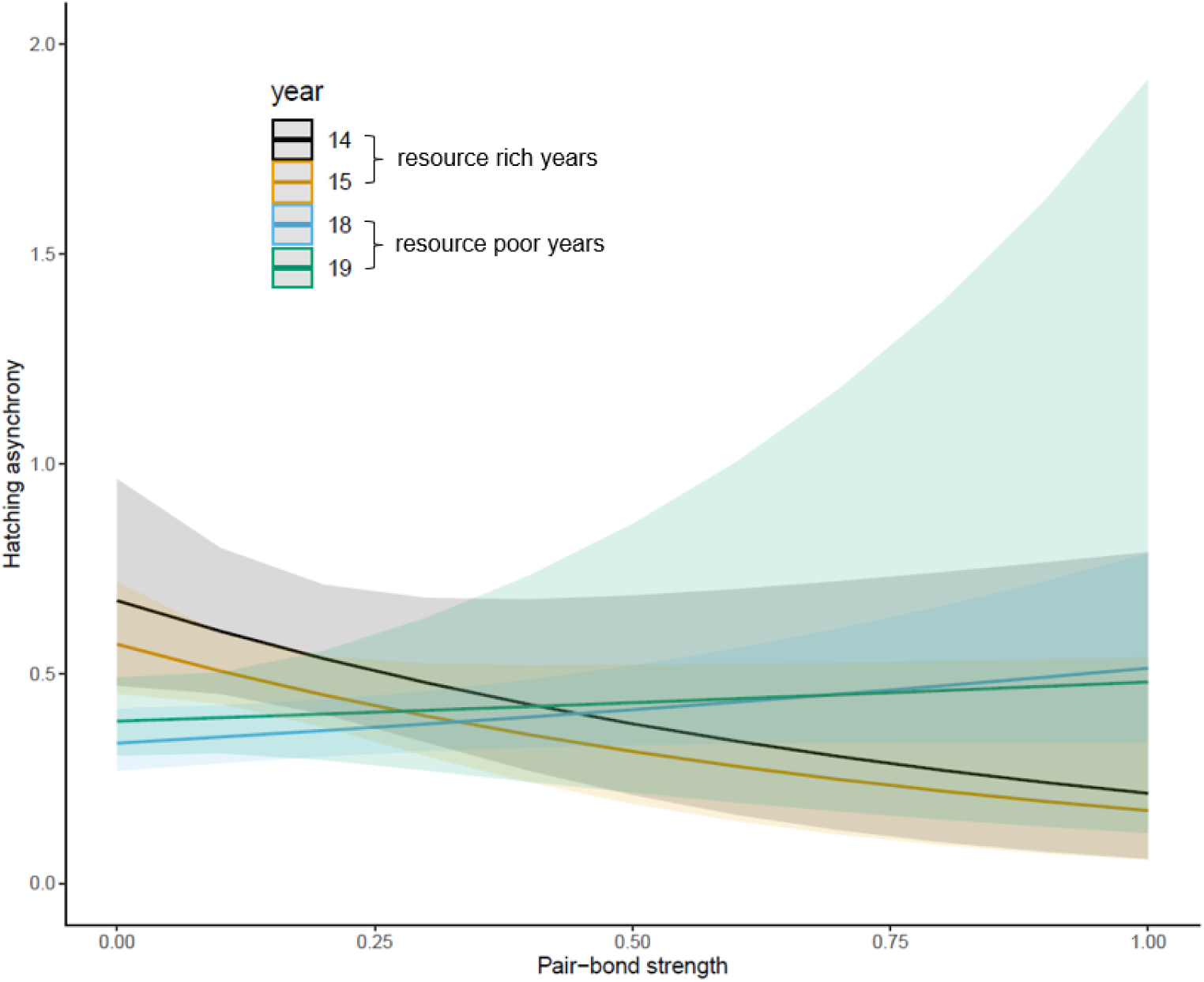
We found a significant interaction between pair-bond strength and year (a proxy of environmental conditions) on hatching synchrony. Here, a higher value on the y-axis represents more asynchronous hatching. Note that for visualisation purposes, pair-bond strength (log-transformed PCA values) is standardised between 0 and 1.

#### (b) Success in fledging offspring

Pair-bond strength was not associated with the number of fledglings, mass of fledglings, proportion of hatched chicks that fledged or cumulative fledging (Table 1; Figure 4). According to population-wide models of number and mass of fledglings per year, 2018 and 2019 were poor years for jackdaw reproductive success relative to 2014 and 2015 (Figure 3; see Supplementary Materials for further details). Pairs in 2018 and 2019 fledged significantly fewer chicks than 2015 (N datapoints = 181; N pairs = 106; 2018 relative to 2015: β = −0.15, SE = 0.07, 95% CI [-0.27, −0.01], P = 0.04; 2019 relative to 2015: β = −0.29, SE = 0.07, 95% CI [-0.43, −0.15], P < 0.01), and in 2019 pairs also fledged significantly fewer chicks than in 2014 and 2018 (N datapoints = 181, N pairs = 106; 2019 relative to 2014: β = −0.32, SE = 0.11, 95% CI [-0.53, - 0.12], P <0.01; 2019 relative to 2018: β = −0.15, SE = 0.06, 95% CI [-0.26, −0.02], P = 0.03). Similarly, in 2019 the cumulative mass of fledglings per pair was significantly lower than in 2014, 2015 and 2018 (N datapoints = 181, N pairs = 106; 2019 relative to 2014: β = −0.46, SE = 0.11, 95% CI [-0.66, −0.25], P <0.01; 2019 relative to 2015: β = −0.28, SE = 0.07, 95% CI [-0.42, −0.14], p <0.01; 2019 relative to 2018: β = −0.18, SE = 0.06, 95% CI [-0.29, −0.06], P <0.01). Together these results show that 2019 was the hardest year for jackdaws in terms of reproductive success, followed by 2018. 2014 and 2015 were comparatively good years. All models without the interaction term between year and hatching synchrony were better than models with its inclusion (where a ‘better’ model has an AIC more than or equal to two less than the competing model; Table S3). This suggests that hatching synchrony did not interact with year to influence reproductive outcome, at least across the four years of our study.

**Figure 4.**
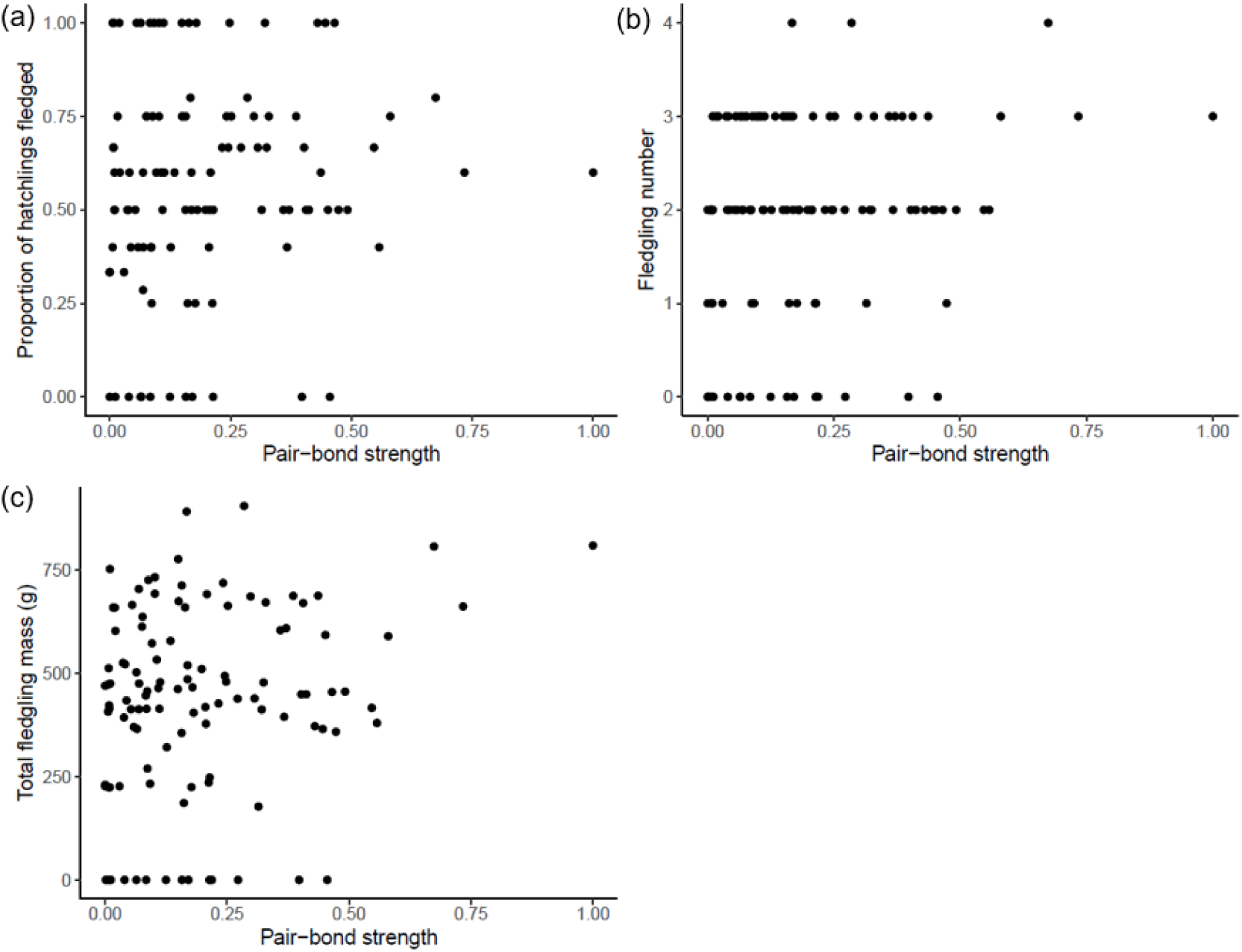
There was no significant relationship between pair-bond strength (mean pair-bond strength per pair per year) and different proxies of reproductive success, such as (a) the proportion of hatchlings that fledged, (b) the number of fledglings, and (c) the total fledgling mass. Note that for visualisation purposes, pair-bond strength (log-transformed PCA values) is standardised between 0 and 1.

**Table 1.** Results of models testing whether pair-bond strength correlated with reproductive success per pair per year, or reproductive success per pair across five years for the response variable ‘cumulative fledgling number’. Pair ID and site were included as random effects for all models except the cumulative fledgling model, where only site was included as a random effect. Full details of variable definitions and additional checks of robustness are given in the Methods. Bold indicates significant results.

| Response variable | n (datapoints, pairs) | Predictor variables | $\beta$ | SE | X2 | 2.5% C I | 97.5% CI | P-value |
| --- | --- | --- | --- | --- | --- | --- | --- | --- |
| Fledgling number | 93,47 | Pair-bond strength | 0.003 | 0.019 | 0.032 | -0.040 | 0.033 | 0.858 |
|  |  | Male age | 0.016 | 0.036 | 0.213 | -0.054 | 0.087 | 0.645 |
|  |  | Year:2015 | -0.133 | 0.116 |  | -0.360 | 0.095 |  |
|  |  | Year:2018 | -0.296 | 0.167 | 3.976 | -0.625 | 0.032 | 0.264 |
|  |  | Year:2019 | -0.384 | 0.194 |  | -0.764 | -0.000 |  |
|  |  | **Food-sharing rate** | **-0.036** | **0.017** | **4.65** | **-0.069** | **0.003** | **0.031** |
|  |  | Male tarsus length | -0.018 | 0.042 | 0.180 | -0.101 | 0.065 | 0.671 |
|  |  | **Female tarsus length** | **0.066** | **0.033** | **4.051** | **0.002** | **0.130** | **0.044** |
|  |  | Lay date | -0.016 | 0.016 | 1.059 | -0.048 | 0.015 | 0.303 |
| Fledgling mass | 95,48 | Pair-bond strength | 0.016 | 0.019 | 0.732 | -0.021 | 0.052 | 0.392 |
|  |  | Male age | -0.004 | 0.032 | 0.012 | -0.067 | 0.060 | 0.911 |
|  |  | Year:2015 | -0.089 | 0.121 |  | -0.326 | 0.148 |  |
|  |  | Year:2018 | -0.230 | 0.162 | 6.545 | -0.547 | 0.087 | 0.088 |
|  |  | Year:2019 | -0.405 | 0.180 |  | -0.758 | -0.052 |  |
|  |  | Food-sharing rate | -0.022 | 0.017 | 1.627 | -0.056 | 0.012 | 0.202 |
|  |  | Male tarsus length | 0.007 | 0.040 | 0.030 | -0.071 | 0.084 | 0.863 |
|  |  | Female tarsus length | 0.048 | 0.031 | 2.409 | -0.013 | 0.108 | 0.121 |
|  |  | Lay date | -0.049 | 0.006 | 2.396 | -0.049 | 0.006 | 0.122 |
| Proportion hatched that fledged | 94,47 | Pair-bond strength | -0.006 | 0.035 | 0.035 | -0.075 | 0.062 | 0.853 |
|  |  | Male age | -0.085 | 0.075 | 1.297 | -0.232 | 0.061 | 0.255 |
|  |  | Year:2015 | -0.017 | 0.252 |  | -0.512 | 0.478 |  |
|  |  | Year:2018 | -0.120 | 0.289 | 1.844 | -0.687 | 0.447 | 0.605 |
|  |  | Year:2019 | -0.362 | 0.342 |  | -1.032 | 0.308 |  |
|  |  | Food-sharing rate | -0.047 | 0.045 | 1.078 | -0.135 | 0.042 | 0.299 |
|  |  | Male tarsus length | -0.027 | 0.072 | 0.143 | -0.168 | 0.113 | 0.705 |
|  |  | Female tarsus length | 0.087 | 0.054 | 2.609 | -0.018 | 0.192 | 0.106 |
|  |  | Lay date | -0.035 | 0.031 | 1.284 | -0.095 | 0.025 | 0.257 |
| Cumulative fledgling | 12,12 | Mean pair-bond strength | 0.113 | 0.140 | 0.650 | -0.161 | 0.387 | 0.420 |
|  |  | Male tarsus length | 0.122 | 0.107 | 1.290 | -0.088 | 0.332 | 0.256 |
|  |  | Female tarsus length | 0.089 | 0.072 | 1.549 | -0.051 | 0.229 | 0.213 |

## Discussion

In this study we examined four key assumptions of the *Social Intelligence Hypothesis (SIH)* in pair-bonding birds. We found evidence that pair-bond strength (1) varied between pairs, (2) was consistent within pairs, and (3) was positively associated with a measure of partner responsiveness. However (4) although pairs with stronger bonds were better able to adjust hatching synchrony to current environmental circumstances, we found no evidence that pair-bond strength was linked associated with reproductive success

### Does pair-bond strength (1) vary between pairs and is it (2) consistent within-pairs?

We found clear evidence that in our jackdaw populations pair-bond strength varied between pairs and was consistent within them. The repeatability estimates of pair-bond strength in our study are higher than the average repeatability of behaviour in general (0.37) as reported in a meta-analysis of behavioural consistency (71), and similar to the repeatability of social network position (0.41 – 0.62) in wild great tits (*Parus major*) (72). These results are consistent with the assumption that the ability to maintain strong bonds represents a stable trait that could come under selection (25). While variation in social bonding could reflect a multiplicity of underlying factors including endocrine profiles (73), individual personality (74), and familiarity (44), the *SIH* assumes a central role for information-processing abilities (25). We turn our attention to this assumption in the next section.

### (3) Is pair-bond strength related to partner responsiveness?

Our results suggest that in more strongly bonded pairs the male was more responsive to his partner. While males responded in variable ways to their partner following the partner’s exposure to a stressor (in both experimental and natural contexts (56)), males in stronger pair-bonds showed a larger absolute change in partner-directed behaviour (contact and allopreening behaviour). Given that male responses were not linked to any overt changes in female behaviour, this indicates that males in stronger bonds are more responsive to very subtle cues of female state. This is reminiscent of findings in humans, where individuals who show better socio-cognitive performance (e.g., better recognition of and response to the emotional state of other individuals) form stronger friendships (75). However, although our findings imply an important role for cognition in the broad sense (11) because males must detect, process and act upon information about their partner’s state (i.e., exhibit social competence (76)), the specific cognitive mechanisms are unknown. One possibility is that responses are driven by emotional contagion, where information about a partner’s state leads to matching states between partners (48). Elucidating the mechanisms underlying social bonding is complicated by the fact that bonds are a product of the behaviour of both partners. Evaluating reciprocal responsiveness by both partners is logistically challenging but will be an important focus for future work. We also note that we cannot unequivocally rule out the possibility that weakly bonded males were just as capable as strongly bonded counterparts of detecting their partner’s state but were simply less motivated to respond. Nevertheless, we would argue that to understand the potential for selection to shape partner responsiveness (and the underlying proximate mechanisms), what matters is how information-processing translates into action.

### (4) Do pairs with stronger bonds have higher fitness?

While our results provide no evidence linking pair-bond strength directly with direct proxies of reproductive success, they do suggest bond strength may be linked to variation in hatching asynchrony. Specifically, relative to weakly-bonded pairs, more strongly bonded pairs hatched their clutches more synchronously in good years and less synchronously in poor years. Thus, the behaviour of strongly bonded pairs appears to match adaptive hypotheses on optimal asynchrony strategies which suggest that more asynchronous broods are favoured in less productive years (66). This suggests that pairs with stronger bonds may have been better able to adjust their hatching synchronicity to environmental conditions. The precise mechanisms through which more strongly bonded pairs may be better at adjusting hatching synchrony to environmental conditions are unclear. Hatching synchrony is, however, related to incubation initiation (70), which is under female control in jackdaws (55). If males do not respond to female incubation initiation cues and do not food-share with their partner, then incubation initiation will be disrupted because the female must leave the nest box in order to feed. Therefore, one explanation as to why more strongly bonded pairs are better able to adjust hatching synchrony to environmental conditions is that males are more responsive to their partner’s behaviour, and thus can better coordinate the initiation of incubation.

In theory, the adjustment of hatching synchrony to environmental conditions should result in fitness benefits (66,67), but we detected no signal that hatching synchrony interacted with year to influence reproductive success across the four years analysed. We also found no effect of pair-bond strength on the total number and mass of fledglings per pair per year, or cumulative fledging success per pairs over multiple years. The absence of a relationship between pair-bond strength and fitness is at odds with previous work showing that pair-bond strength correlates positively with fledging success in captive cockatiels (45), and that pair-bond duration, the stability of the bond and the familiarity of partners (likely to be facets of pair-bond strength) influence fitness outcomes (42,44,77–80). Moreover, this result could be seen to undermine a central tenet of the *SIH*, suggesting that forming and maintaining strong relationships may not be a key driver of cognitive evolution in large-brained birds (cf. (25,35)).

Before drawing this conclusion, however, alternative hypotheses must be addressed. First, survival is a key component of fitness that we were not able to test in this study. In multiple species, social bonds have been linked to increased probability of survival for adults (7,81,82) and their offspring (6,83,84). Investigating whether individuals in strong pair-bonds have a higher probability of survival, or if their offspring have a higher probability of survival post-fledging, is a vital step forward in elucidating whether pair-bond strength influences fitness. Second, while the pair-bond is the most valuable relationship in corvid society, pairs do not exist in a social vacuum but are embedded within wider social networks and navigate numerous other social bonds (25,85). For instance, partners work together to interrupt relationship formation between potential competitors in ravens (*Corvus corax)* (86) while in jackdaws and rooks (*Corvus frugilegus*) partners aid each other in fights against third parties (87) and learn from and associate with flock-members independently of one another (52,88). There is also some evidence that jackdaws with more central social network positions have better reproductive outcomes (89). Given that time spent together is an important component of pair-bond strength, and that it takes time to monitor, form and maintain non-pair relationships, there is an implied trade-off between the management and maintenance of pair versus non-pair relationships. Indeed, in humans there is evidence of a trade-off between the quality of a relationship with a romantic partner and the quantity of non-romantic social bonds (90,91), while in jackdaws we have found that investment in pair-bonds trades off against the strategic adjustment of social associations in the wider social network (8). Such trade-offs could limit strongly bonded partners’ access to valuable social and cultural information and so obscure the detection of a direct relationship between pair-bond strength and reproductive success. Studies examining how animals navigate the potentially conflicting demands of different types of relationship are now needed to characterise such trade-offs and deepen our understanding of the cognitive demands of sociality. Finally, while we have treated bond strength as a trait that may impact fitness, in reality the strength of social bonds is an emergent product of two (or more) interacting individuals. Future research will therefore need to consider the impacts of interacting genotypes (indirect genetic effects; (92)) to fully understand how selection shapes social bonding and the underlying cognitive processes.

## Conclusion

The *Social Intelligence Hypothesis* (and its avian-specific formulation, the *Relationship Intelligence Hypothesis*) emphasise long-term, cooperative social bonds as central to understanding cognitive and brain evolution. We found support for three key assumptions of the *SIH* in wild corvids, birds renowned for their sophisticated cognitive abilities and large brains: the strength of relationships varies between mated pairs of jackdaws, is consistent within pairs and is linked to measures of responsiveness to partner states. However, while pairs with stronger bonds may be better able to co-ordinate reproductive behaviour in response to variable environmental conditions, we did not find any evidence that pair-bond strength influences reproductive success. In a field largely dominated by broad-scale comparative studies, further intra-specific testing of links between cognitive performance, social behaviour and fitness are vital to resolve debates as to whether and how social relationships may drive cognitive evolution. We propose that studies investigating how animals navigate trade-offs between long-term partnerships and wider social connections are now vital to understand the cognitive demands of social life and their evolutionary consequences.

## Ethics

This study was carried out with approval from the University of Exeter Biosciences Research Ethics Committee (eCORN002970; eCORN001858) following the ASAB Guidelines for the Treatment of Animals in Behavioural Research and Teaching (93). All subjects were ringed, and blood was collected by Cornish Jackdaw Project team members licensed by the British Trust for Ornithology and UK Home Office (project licence 30/3261).

## Data availability

Data and scripts are available at https://figshare.com/s/23cb3f83ee8c9fd999bf.

## Acknowledgements

We are grateful to the Gluyas family and Odette Eddy for allowing us access to their land to conduct this research. We are also grateful to Ella Meekins, Angélica Bas Gomez, Anna Bowland, Amy Hall, Coby Thompson-Knight, Emma Doyle, Emily Cuff, Lucy Penney, Gray Wirtanen, and Sam Mosedale for their help with coding video data. Thank you also to Neeltje Boogert and Josh Arbon for providing helpful feedback on the manuscript.

## Supplementary methods

### Supplementary Tables

**Table S1.** The behavioural ethogram used in BORIS v7.4.6 to code video data.

| Behaviour | Type | Description |
| --- | --- | --- |
| **ALLOPREEN** | Duration | One individual preens another (Figure 1a) |
| **CONTACT** | Duration | An individual is stood or sat close enough to their partner that they would not have to move their bodies in order to make physical contact (i.e., they are within a beak's distance of one another). They are not actively engaged in any other behaviour. |
| **CHATTER** | Duration | A specific call often made between partners. |
| **IN** | Duration | An individual is in the box (time together and male visit rate are extracted from this variable). |
| **FS** | Point | Food-sharing between adults. In the incubation stage this is always from the male to the female. Coded only once per male visit to the box. |
| **PEEK (vigilance)** | Duration | Individual looks out of nest-box. |
| **LAY (incubation)** | Duration | Individual sitting on eggs (start behaviour when the individual 'wiggles' onto the eggs). |
| **COPULATION** | Duration | One individual attempt to or does copulate with another. |

**Table S2.**
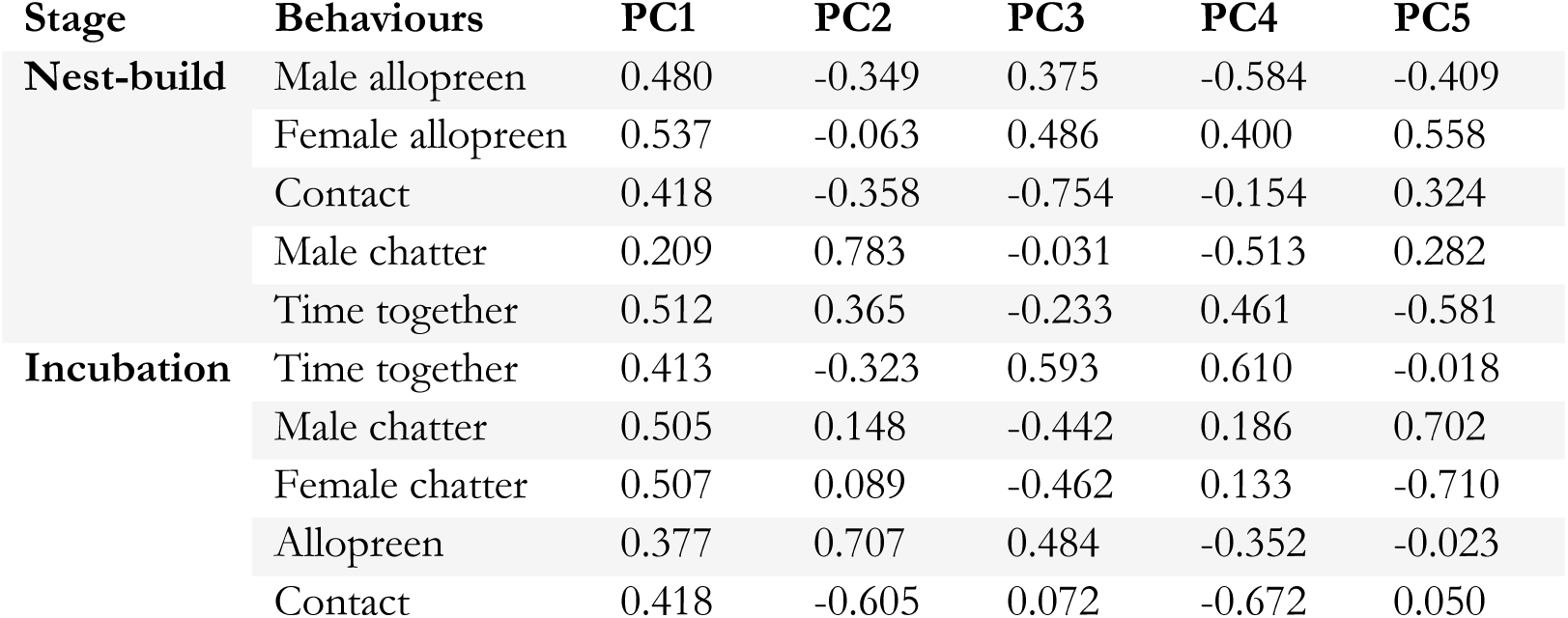
PCA loadings for the nest-building and incubation stage.

| Stage | Behaviours | PC1 | PC2 | PC3 | PC4 | PC5 |
| --- | --- | --- | --- | --- | --- | --- |
| **Nest-build** | Male allopreen | 0.480 | -0.349 | 0.375 | -0.584 | -0.409 |
|  | Female allopreen | 0.537 | -0.063 | 0.486 | 0.400 | 0.558 |
|  | Contact | 0.418 | -0.358 | -0.754 | -0.154 | 0.324 |
|  | Male chatter | 0.209 | 0.783 | -0.031 | -0.513 | 0.282 |
|  | Time together | 0.512 | 0.365 | -0.233 | 0.461 | -0.581 |
| **Incubation** | Time together | 0.413 | -0.323 | 0.593 | 0.610 | -0.018 |
|  | Male chatter | 0.505 | 0.148 | -0.442 | 0.186 | 0.702 |
|  | Female chatter | 0.507 | 0.089 | -0.462 | 0.133 | -0.710 |
|  | Allopreen | 0.377 | 0.707 | 0.484 | -0.352 | -0.023 |
|  | Contact | 0.418 | -0.605 | 0.072 | -0.672 | 0.050 |

**Table S3.**
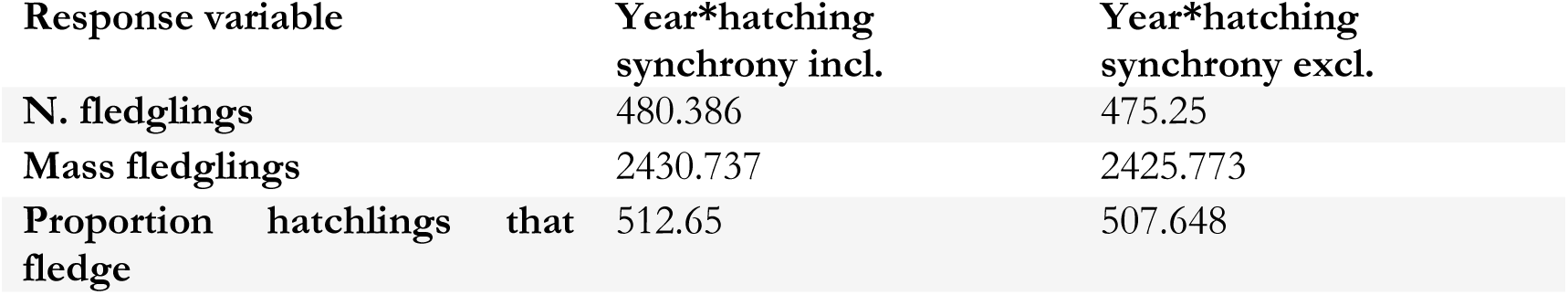
AIC values for models including the interaction between year and hatching synchrony and excluding it. Models excluding the interaction term always performed better.

| Response variable | Year*hatching synchrony incl. | Year*hatching synchrony excl. |
| --- | --- | --- |
| N. fledglings | 480.386 | 475.25 |
| Mass fledglings | 2430.737 | 2425.773 |
| Proportion hatchlings that fledge | 512.65 | 507.648 |

### Supplementary Figures

**Figure S1.**
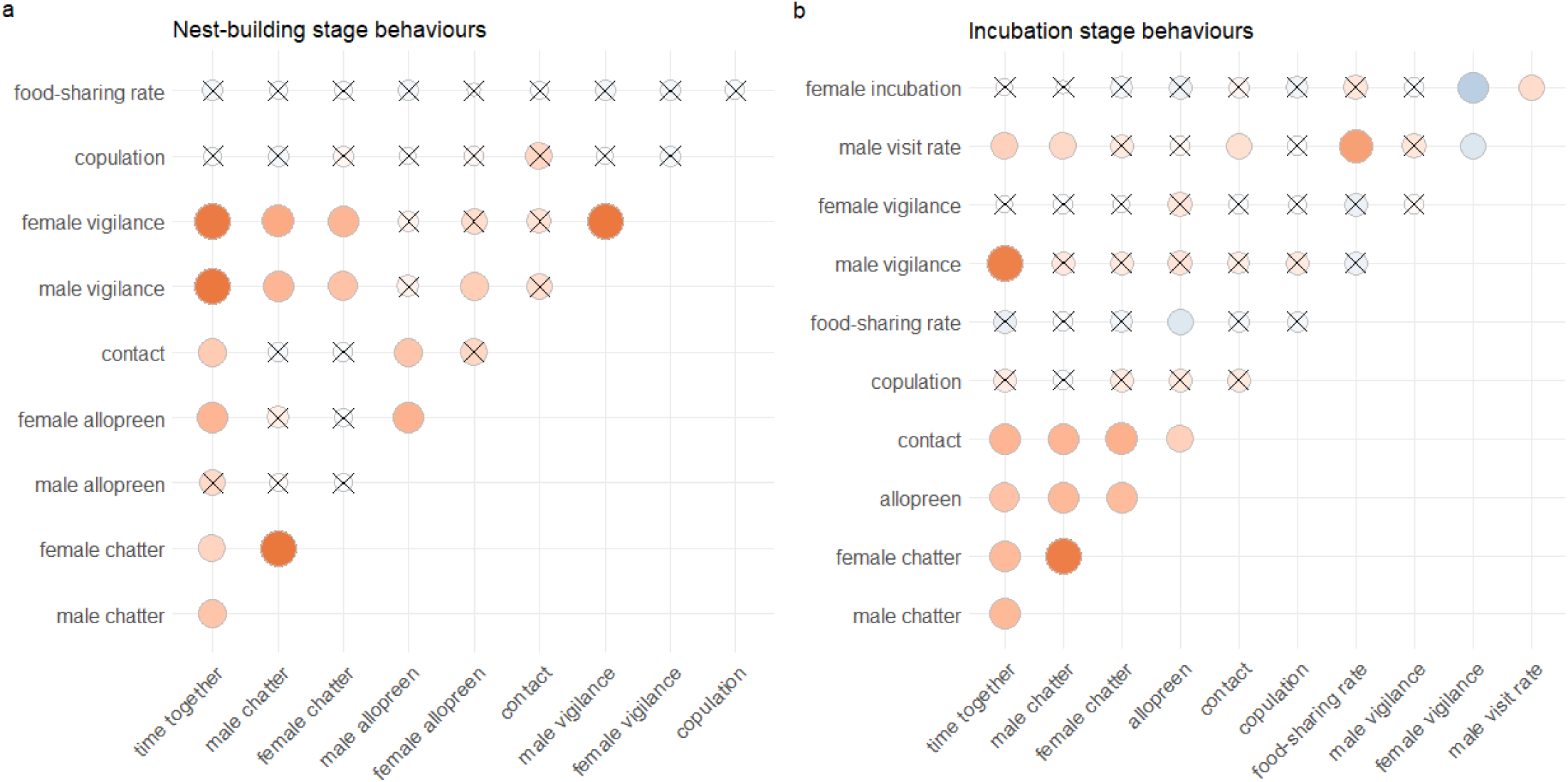
Correlation plots of behaviours in (a) the nest-building stage of the breeding season and (b) the incubation stage of the breeding season. Crosses indicate a non-significant (α > 0.05) pairwise correlation. Red indicates a positive correlation; blue indicates a negative correlation.

**Figure S2.**
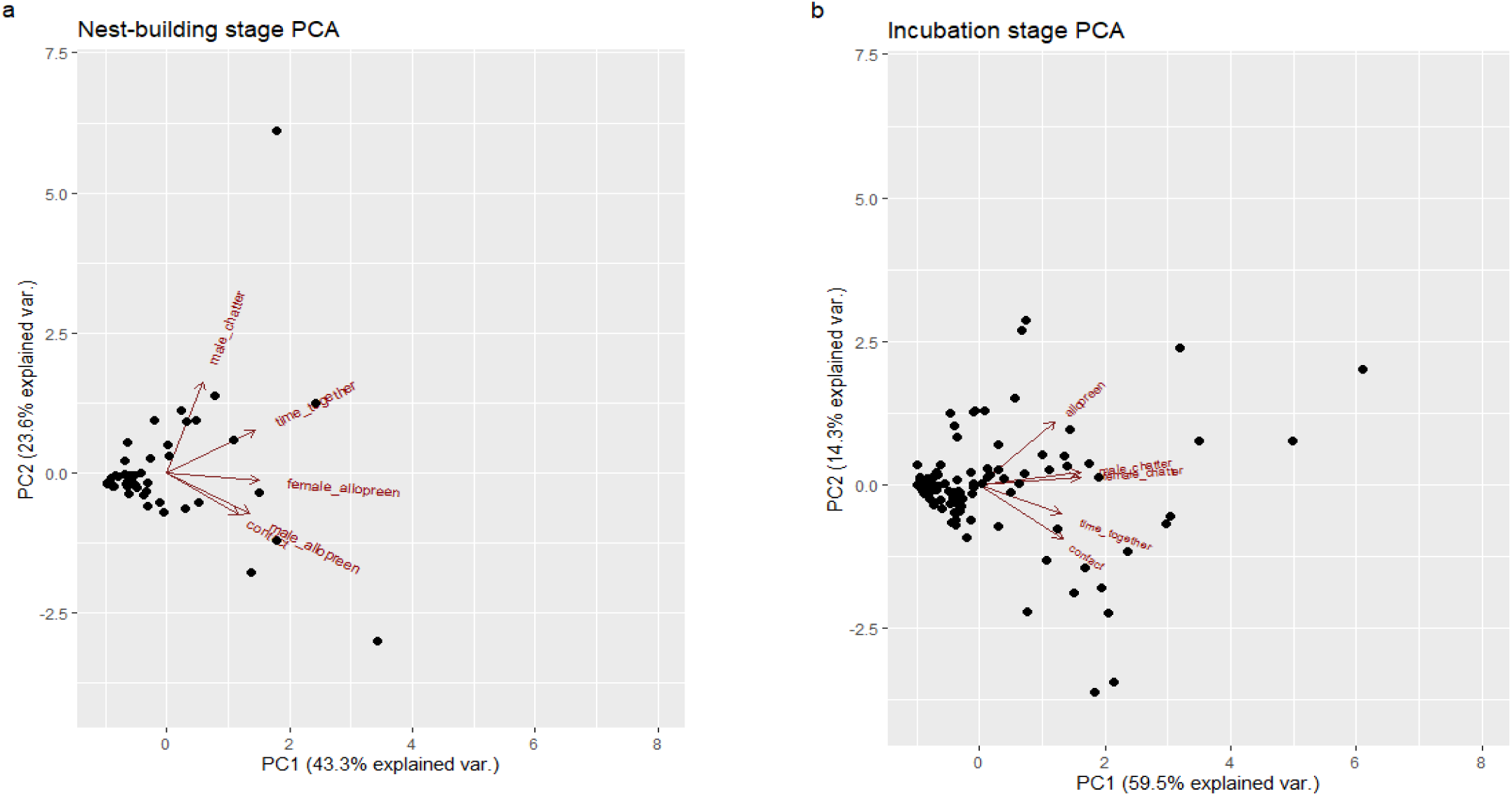
PCA biplots of (a) nest-building and (b) incubation stage affiliative behaviours. Note that for b, the x axis has been truncated to 8 so that loadings are visible; one datapoint at (13,0.5) is not visible.

**Figure S3.**
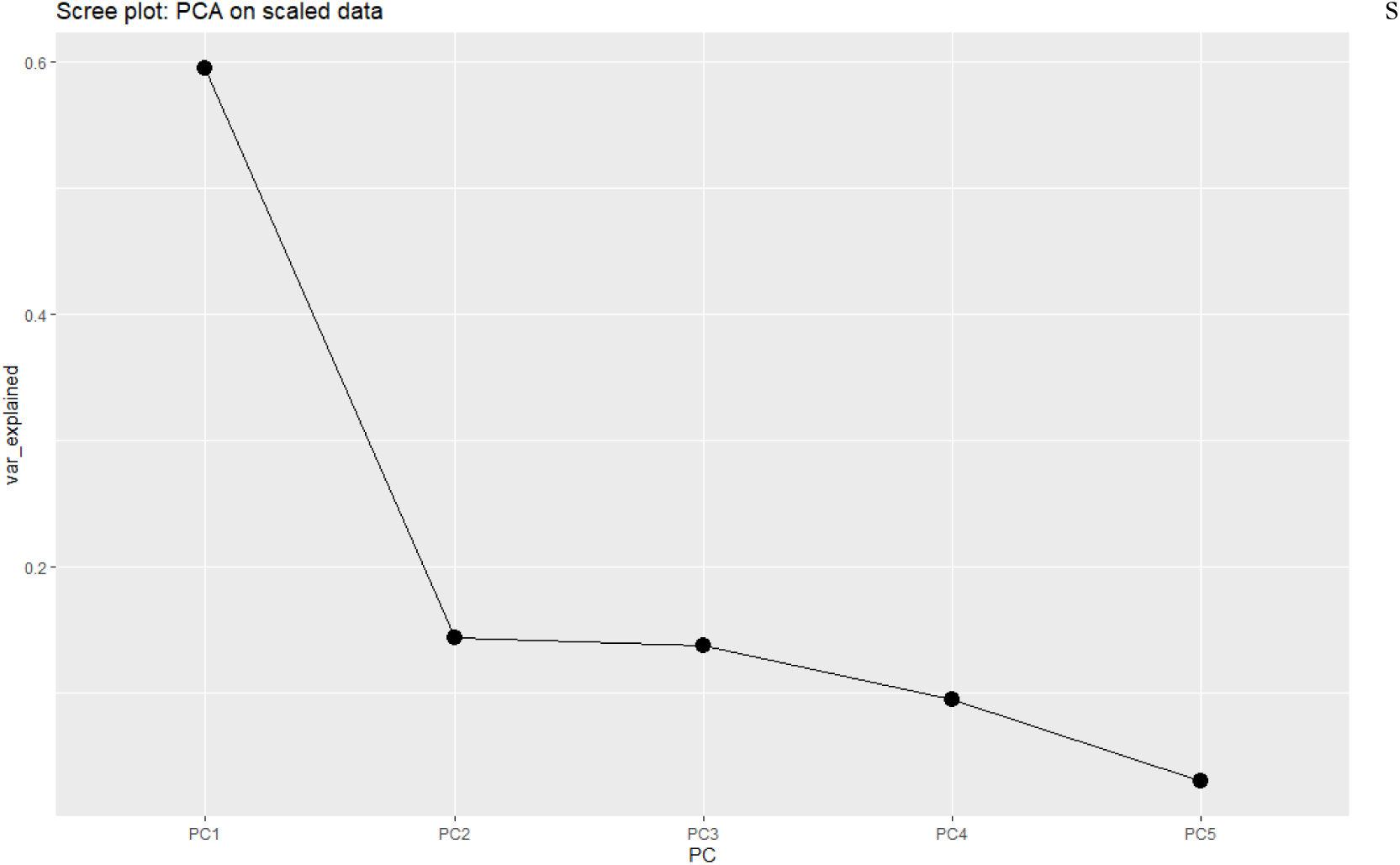
A scree plot showing variation explained by each Principal Component in the incubation-stage PCA. Although affiliative behaviours loaded onto both PC1 and PC2, the scree plot shows clear justification for keeping only PC1 for further analyses given the relative amount of variation explained.

